# Metabarcoding reveals different zooplankton communities in northern and southern areas of the North Sea

**DOI:** 10.1101/2020.07.23.218479

**Authors:** Jan Niklas Macher, Berry B. van der Hoorn, Katja T. C. A. Peijnenburg, Lodewijk van Walraven, Willem Renema

## Abstract

Zooplankton are key players in marine ecosystems, linking primary production to higher trophic levels. The high abundance and high taxonomic diversity renders zooplankton ideal for biodiversity monitoring. However, taxonomic identification of the zooplankton assemblage is challenging due to its high diversity, subtle morphological differences and the presence of many meroplanktonic species, especially in coastal seas. Molecular techniques such as metabarcoding can help with rapid processing and identification of taxa in complex samples, and are therefore promising tools for identifying zooplankton communities. In this study, we applied metabarcoding of the mitochondrial cytochrome c oxidase I gene to zooplankton samples collected along a latitudinal transect in the North Sea, a shelf sea of the Atlantic Ocean. Northern regions of the North Sea are influenced by inflow of oceanic Atlantic waters, whereas the southern parts are characterised by more coastal waters. Our metabarcoding results indicated strong differences in zooplankton community composition between northern and southern areas of the North Sea, particularly in the classes Copepoda, Actinopterygii (ray-finned fishes) and Polychaeta. We compared these results to the known distributions of species reported in previous studies, and by comparing the abundance of copepods to data obtained from the Continuous Plankton Recorder (CPR). We found that our metabarcoding results are mostly congruent with the reported distribution and abundance patterns of zooplankton species in the North Sea. Our results highlight the power of metabarcoding to rapidly assess complex zooplankton samples, and we suggest that the technique could be used in future monitoring campaigns and biodiversity assessments.

**Highlights:**

- Zooplankton communities are different in northern and southern areas of the North Sea
- Metabarcoding results are consistent with known species distributions and abundance
- Metabarcoding allows for fast identification of meroplanktonic species

## Introduction

Zooplankton are key players in marine ecosystems, linking primary production to higher trophic levels (Suthers & Rissik, 2009; Beaugrand, Edwards & Legendre, 2010; Turner, 2015; Steinberg & Landry, 2017). Due to their abundance and high taxonomic diversity, zooplankton can be used for ecosystem assessments and biomonitoring (Bucklin et al., 2016; Chain et al., 2016; Chiba et al., 2018). However, studying zooplankton is a challenging task, as obtaining samples and taxonomic identification can be difficult (Schminke, 2007; McManus & Katz, 2009; Cornils & Held, 2014). Especially meroplanktonic species, i.e. taxa which are part of the plankton only during their larval stages, can be difficult to identify (Mathivat-Lallier & Cazaux, 1990; Kirby & Lindley, 2005; Oozeki, 2018). This is also reflected in the extensive, publicly available long-term monitoring dataset of the CPR (Continuous Plankton Recorder, available online: https://data.cprsurvey.org/datacatalog; (Reid et al., 2003a) which records the occurrence of more than 30 copepod taxa on genus or species level, whereas data on meroplanktonic groups is available in less detail (e.g.“fish eggs”, “polychaete larvae”). Inclusion of the often highly abundant meroplanktonic species, especially in coastal areas (Schwamborn et al., 2001; Kirby & Lindley, 2005; Jansen et al., 2012; Harvey et al., 2018) in zooplankton biodiversity assessments would be beneficial for getting more detailed insights and for better understanding of zooplankton distribution patterns. Molecular techniques like metabarcoding (Taberlet et al., 2012), i.e. the amplification, sequencing and analysis of marker gene fragments (“molecular barcodes”, (Ratnasingham & Hebert, 2007)) of whole communities, can help with rapid processing and identification of species in complex samples. The technique has been shown to be an effective tool for identification of species in zooplankton communities (Brown et al., 2015; Casas, Pearman & Irigoien, 2017; Deagle et al., 2018; Zhang et al., 2018) and for identification of larval stages (Kimmerling et al., 2018; Couton et al., 2019). While several studies have shown the benefits of metabarcoding zooplankton, suitable barcoding regions and primers for amplification are still under discussion (Brown et al., 2015; Bucklin et al., 2016; Chain et al., 2016; Clarke et al., 2017), and current DNA reference databases are far from complete (Bucklin et al., 2016). However, the development of highly degenerate primers amplifying a wide range of taxa is an important step towards assessment of complex communities (Leray et al., 2013; Wangensteen et al., 2018) and is therefore especially promising for the assessment of highly diverse zooplankton communities.

In this study we use the highly degenerate Leray XT primers (Leray et al., 2013; Wangensteen et al., 2018), which amplify a fragment of the mitochondrial cytochrome c oxidase I gene, to assess the zooplankton community of the North Sea along a transect from the Dutch coast to the Shetland Islands. The zooplankton of the North Sea, a shelf sea of the Atlantic Ocean, is relatively well known based on morphological analyses (Fransz et al., 1991; Greve et al., 2001; Beare et al., 2002; Lindley & Batten, 2002; Reid et al., 2003b; Alvarez-Fernandez, Lindeboom & Meesters, 2012). Previous studies have shown that the zooplankton community in the northern parts of the North Sea shows a higher abundance of oceanic species, while the community in the southern parts of the North Sea is commonly dominated by more coastal species (Fransz et al., 1991; Krause, M., Dippner, J.W., Beil, J., 1995; Nielsen & Sabatini, 1996; Alvarez-Fernandez, Lindeboom & Meesters, 2012). The community structure is linked to the influx of cold, saline, Atlantic waters entering the North Sea from the north, and flowing south through a corridor of deeper water to the area of the Dogger Bank in the central North Sea (Otto et al., 1990; Fransz et al., 1991; Lindley & Batten, 2002). We hypothesised that metabarcoding of the zooplankton across a latitudinal transect of the North Sea, using highly degenerate COI primers, would allow for the identification of distinct zooplankton communities in the northern and southern parts of the North Sea.

## Material & Methods

### Sampling, DNA extraction and library preparation

Samples from nine stations were taken during the North Sea leg 10 of the NICO (Netherlands Initiative Changing Oceans) expedition in May and June 2018 (see supplementary Table S1 for coordinates and Figure 1 for a map). Sampling in UK waters was approved by the Maritime Policy Unit (Legal Directorate) of the Foreign and Commonwealth Office (ref 33/2018). Weather conditions were calm and stable throughout the 12 day cruise. Samples were taken with a plankton MultiNet (Hydro-Bios, Kiel, Germany) with a mesh size of 100 μm. All tows were conducted between 8:00 and 9:00 in the morning. Winch speed was 5 meters per second. Temperature and salinity across the water column were measured by the on-board CTD prior to sampling. Zooplankton samples were taken from the seafloor to the deepest thermocline, between thermoclines (in case more than one was present), and from the uppermost thermocline to the surface. Zooplankton was removed from the multinet by carefully rinsing the net with seawater into the cod end. Samples were transferred to a Folsom sample splitter (McEwen, Johnson & Folsom, 1954). Half of the sample volume was retained for morphological analyses and half for molecular analyses. The molecular subsample was further split in two halves, which were subsequently processed as separate samples (i.e. extraction replicates) to check for potential biases during processing and sequencing. All samples were transferred to 96% ethanol and stored at −20°C until further processing.

**Figure 1:**
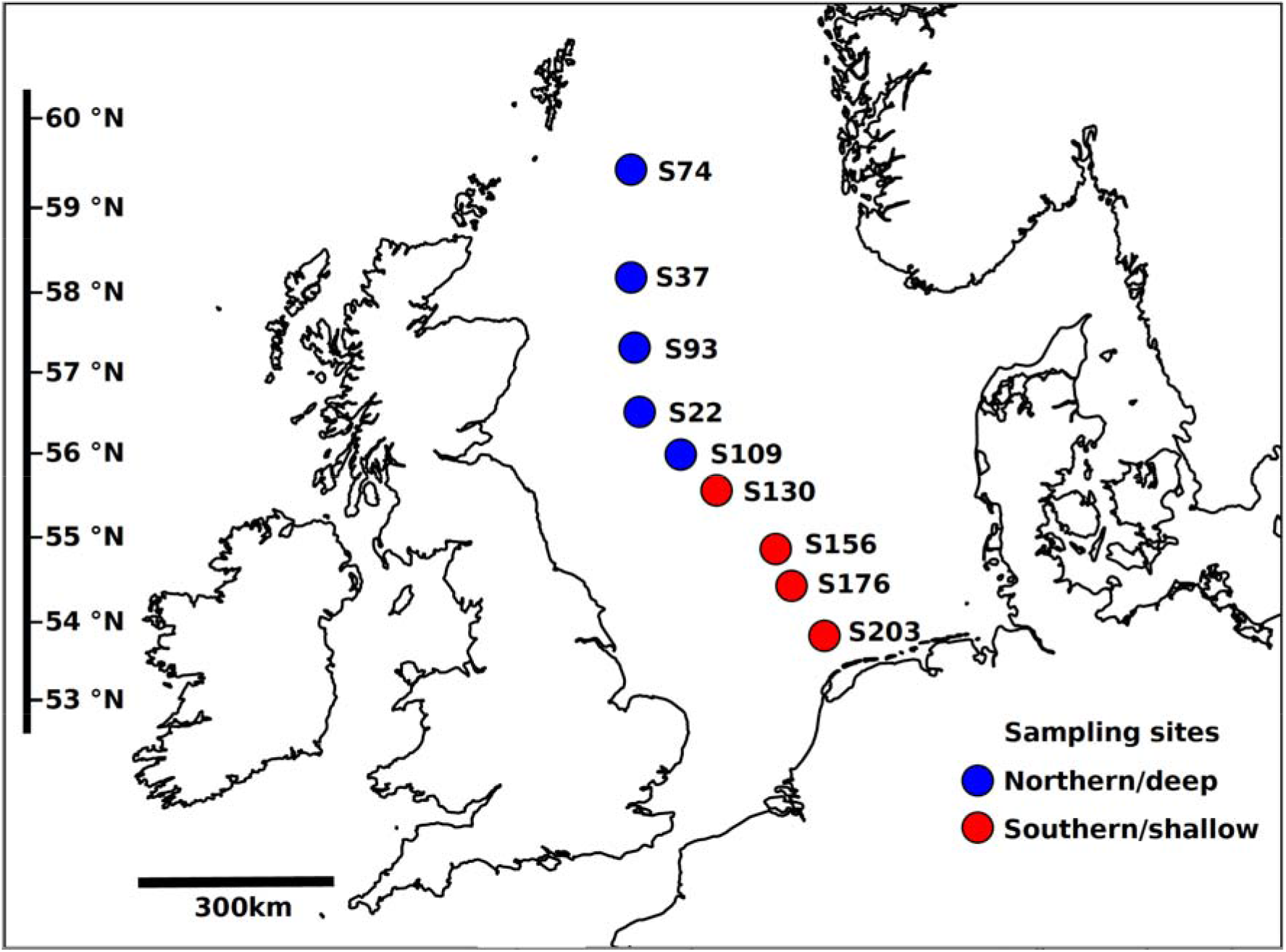
Map showing the location and station names of the nine sampling sites in the North Sea. Blue circles and red circles represent ‘northern/deep’ and ‘southern/shallow’ sampling sites, respectively.

Samples were dried under sterile fume hoods and ground to a fine powder using an IKA Ultra Turrax homogenizer (IKA, Staufen, Germany) on full speed for 10 minutes (Macher et al., 2018; Zizka et al., 2019b). DNA was extracted using the Macherey-Nagel (Düren, Germany) NucleoSpin tissue kit on the KingFisher (Waltham, USA) robotic platform, following the manufacturer’s protocol. Two negative controls containing ultrapure water were processed together with the samples during all steps. Quantity and size of the extracted DNA was checked on the QIAxcel platform (Qiagen, Hilden, Germany). 15ng of DNA per sample was used for metabarcoding using the Leray-XT primers (313bp product length), which amplify a COI gene fragment of a wide range of marine metazoan taxa (Wangensteen et al., 2018). Samples were amplified using a two-step PCR protocol as commonly used for metabarcoding studies (Andruszkiewicz et al., 2017; Galan et al., 2018; Zizka et al., 2019a). The first PCR was performed in 20 μl PCR reactions containing 10 μl Environmental Master Mix (2x, Thermo Fisher Scientific, Waltham, USA), 7μl ultrapure water, 1ul of each primer (10pMol/ul) and 1 μl (15ng) of DNA template. PCR was conducted with 10 minutes of initial denaturation at 95°C, followed by 30 cycles of 30 seconds denaturation at 95°C, 30 seconds annealing at 50°C, and 20 seconds extension at 72°C. Final extension was set to 7 minutes at 72°C. Amplicons were cleaned with Macherey Nagel NucleoMag beads (Dueren, Germany) according to the manufacturer’s protocol and a sample to beads volume of 1:0.9.

The second PCR step was used to tag samples with unique Illumina adapters. Samples were amplified using 7ul of ultrapure water, 10ul of Environmental Master Mix, 1 μl of each primer tagged with Nextera XT adapter (Illumina, San Diego, USA) and 1ul of DNA template. Cycling conditions were the same as described above, but only 10 cycles were used. Amplicon length and concentration was measured on the QIAxcel platform, samples were cleaned and size selected using magnetic beads as described above, and equimolarly pooled using the QIAgility platform (Qiagen, Venlo, Netherlands). Negative controls did not show DNA and were added to the library with 10% of the final volume. Final concentration and fragment length of the library were checked on the Bioanalyzer platform (Agilent Technologies, Santa Clara, USA). The final library was sent for sequencing on the Illumina MiSeq platform (2×300bp read length) at Baseclear (Leiden, Netherlands).

### Bioinformatic processing

Processing of reads was conducted using the Galaxy platform (Afgan et al., 2018) following the principal steps of (Beentjes et al., 2019). Samples taken from different depths of the same sampling station were combined to allow for studying the zooplankton community of the entire watercolumn. FLASH (Magoč & Salzberg, 2011) was used to merge reads with minimum overlap of 50 and maximum overlap of 300, a maximum mismatch ratio of 0.2, and with non-merged reads discarded. Cutadapt was used to trim primers (settings: both primers need to be present, minimum number of matching bases 10, maximum error rate 0.2). PrinSeq (Schmieder & Edwards, 2011) was used to filter and trim sequences to 310 base pairs to remove reads that contain gaps or indels, which can be present due to amplification of non-eukaryotic taxa (Wangensteen et al., 2018; Macher et al., 2018; Collins et al., 2019). UNOISE (Edgar, 2016) was used for clustering of Operational Taxonomic Units (OTUs). We chose thresholds of alpha = 4 and a minimum number of 10 reads for the denoising approach, which is similar to settings reported in previous studies that found an alpha of 5 to give reliable results (Elbrecht et al., 2018; Turon et al., 2019). We chose an alpha of 4 to be slightly more restrictive and remove more potentially wrong sequence variants from the dataset, although this approach might also increase the loss of genuine variants. To further reduce the risk of analysing spurious OTUs, only those OTUs with >0.002% relative abundance per sample were retained, which corresponds to >1 read in the sample with the lowest read count. Further, we only retained OTUs that were present in both extraction replicates per sample. Such an approach, i.e. filtering out low abundant OTUs based on relative abundance, is commonly used in metabarcoding studies (Elbrecht et al., 2017; Pereira-da-Conceicoa et al., 2019; Theissinger et al., 2019). After this quality filtering step, the reads of the two technical replicates per sample were summed up to build the final dataset. Quality filtered reads were assigned to species using the BOLD database (Ratnasingham & Hebert, 2007) with the BOLDigger tool (Buchner & Leese, 2020). The following identity thresholds were used for assigning taxonomic ranks: species 98%; genus 95%; family 93%; order 90%; class 85%. OTUs that were assigned to the same taxonomic name were subsequently lumped by summing up reads to prevent analyses of intraspecific variability as provided by the UNOISE pipeline. We focussed our analyses on planktonic metazoans (animal zooplankton).

### Community analyses

We tested which of the parameters: salinity, temperature, bottom depth or latitude, best explained the community composition of zooplankton in the North Sea during the NICO 10 expedition. Analyses of community composition were conducted using the R package vegan (Oksanen et al. 2019, https://cran.r-project.org/package=vegan). Averages of the abiotic variables ‘salinity’ and ‘temperature’ across the water column were obtained from the CTD data (Supplementary Table 1). The variables ‘bottom depth at sampling site’ and ‘latitude of sampling site’ were extracted from the ship logbook. The variables were categorized into two classes (<50th percentile, ≥ 50th percentile of the variable range), and sampling sites were assigned to these classes accordingly. The four southernmost sampling sites (south of 56°N) were also the shallowest (shallower than 75m), while the five northern sampling sites (north of 56°N) were all deeper than 75m. Sites were therefore categorized as ‘southern/shallow’ and ‘northern/deep’, respectively (Fig. 1). Mean salinity was lowest in the three southernmost sampling sites. Mean water temperature was lowest in the sampling sites S130 (55.62 °N), S93 (57.36 °N), and S74 (59.42 °N), i.e. did not show a clear latitudinal pattern. Differences in community composition of the entire zooplankton assemblage as a function of the tested variables were analysed based on relative abundance, i.e. read counts transformed to relative abundance per sample. These analyses were conducted on the level of molecularly identified species. The abundant and species-rich classes Actinopterygii (ray-finned fishes), Copepoda and Polychaeta were also analysed separately to test whether similar patterns could be observed for different taxonomic groups.

Bray- Curtis distances were calculated using the vegdist function implemented in the vegan package. Communities were subsequently clustered with an average-linkage algorithm (hclust function) as in (Burdon et al., 2016; Macher et al., 2018). Community composition was analysed using the ‘adonis’ PERMANOVA function as implemented in vegan. Analyses were run separately with the abiotic variables (depth, latitude, mean salinity, mean temperature) as predictor and the Bray- Curtis distances as response variables. Following (Nakagawa & Cuthill, 2007) and (Cohen, 2013), we regarded significant results with R^2^ > 0.09 (equivalent to r = 0.30) as moderate, and R^2^ > 0.25 (r = 0.50) as strong. Species numbers found exclusively in ‘northern/deep’ or ‘southern/shallow’ sampling sites, or in both areas were visualised using the Venn diagram creator (available online: https://bioinformatics.psb.ugent.be/webtools/Venn/). Correlation of latitude and relative abundance of species was tested with Pearson correlation analyses using the R package ‘ggpubr’ (Kassambra 2019, https://cran.r-project.org/package=ggpubr). This analysis was conducted for the four most abundant species in the classes Copepoda, Actinopterygii and Polychaeta. For comparison of the copepod metabarcoding data with long-term monitoring data based on morphological identification, the May and June data from 2010 to 2017 (latest available data) of the Continuous Plankton Recorder (CPR) dataset (DOI: 10.17031/1628#year=2010-2017;month=5-6) was used. The CPR data was reduced to the 277 samples in the area between 0.5°E and 4.5°E and 53.5°N and 59.5°N. This corresponds to the area covered during the NICO leg 10 expedition. We compared the metabarcoding data (relative read abundance) with data from the CPR (abundance/m^3^). For the ray-finned fishes and polychaetes using CPR data was not possible, as these taxa are recorded as larvae or eggs without further taxonomic identification. *Oithona similis* in the metabarcoding dataset was compared to the *Oithona* spp. data from the CPR, as *Oithona similis* is not specifically recorded by the CPR, but is by far the most common *Oithona* species in the North Sea (Fransz et al., 1991).

## Results

Zooplankton communities from nine sampling sites across the North Sea were analyzed, and 42,798,930 raw reads were obtained. The two negative controls contained a total of 1204 reads (0.0028% of all reads). As Illumina platforms commonly show a low percentage of tag switching during sequencing (Schnell, Bohmann & Gilbert, 2015) and no DNA was observed in the negative controls during library preparation, no contamination was suspected. After merging of forward and reverse reads and quality filtering, 18,904,404 sequences were retained. Bray- Curtis dissimilarity between extraction replicates of the same sample was low (mean 0.018, standard error of the mean 0.002), and therefore no systematic problem with extraction or laboratory processing was suspected.

### Community composition

A total of 3315 OTUs were obtained. These belonged to 33 taxonomic classes, to which 96.1% of quality filtered reads could be assigned. Of the 33 classes, 26 were identified as animals, while 7 classes (with 0.97% of all reads) were identified as plants and bacteria, and were removed prior to further analysis. The zooplankton classes with the highest abundance (based on read counts) were Copepoda (30.2% of reads, 28 identified species), Actinopterygii (ray-finned fishes; 26.3% of reads, 16 identified species), Sagittoidea (arrow worms, 19.9% of reads, 3 identified species), Branchiopoda (10.1% of reads, 3 identified species), Polychaeta (5.8% of reads, 29 identified species), and Echinoidea (5.7% of reads, 5 identified species). All other classes were present with less than 1% of reads. The 26 classes were assigned to 59 orders, 103 families, 119 genera, and 127 species, to which 75% of all reads could be assigned (see supplementary table S2 for a species list). Community composition of the entire zooplankton assemblage differed significantly and strongly between ‘northern/deep’ and ‘southern/shallow’ sampling sites (R2 = 0.35, p = 0.004) as well as between northern and southern sites categorized based on salinity (R2 = 0.31, p = 0.018, Table 1). Water temperature did not explain overall community composition. Similar results were obtained when focussing only on the copepods and polychaetes. For the ray-finned fishes, ‘salinity/latitude’ best explained community composition (Table 1). Copepods, ray-finned fishes and polychaetes as the most abundant and species-rich groups are discussed in more detail below. As latitude/depth best explained community composition, we further focus on this factor.

**Table 1:** Difference in community composition based on Bray- Curtis distance for all species, copepods, ray-finned fishes and polychaetes. Results show the difference in community composition of ‘northern/deep’ versus ‘southern/shallow’ sampling sites; sites with higher versus lower salinity; and sites with higher versus lower water temperature. R^2^ and p value based on ‘adonis’ analysis.

| Taxon | Latitude/depth | Salinity | Temperature |
| --- | --- | --- | --- |
| | $R^2$ and p value | | |
| **All species** | $R^2 = 0.35$<br>$p = 0.004^{**}$ | $R^2 = 0.31$<br>$p = 0.018^*$ | $R^2 = 0.11$<br>$p = 0.596$ |
| **Copepods** | $R^2 = 0.48$<br>$p = 0.005^{**}$ | $R^2 = 0.4$<br>$p = 0.011^*$ | $R^2 = 0.11$<br>$p = 0.359$ |
| **Ray-finned fishes** | $R^2 = 0.3$<br>$p = 0.028^*$ | $R^2 = 0.37$<br>$p = 0.015^*$ | $R^2 = 0.1$<br>$p = 0.602$ |
| **Polychaetes** | $R^2 = 0.58$<br>$p = 0.008^*$ | $R^2 = 0.41$<br>$p = 0.024^*$ | $R^2 = 0.08$<br>$p = 0.536$ |

Community composition was markedly different between ‘northern/deep’ and ‘southern/shallow’ sampling sites. We found 51 (39.8%) of all identified species in both ‘northern/deep’ and ‘southern/shallow’ sites, 48 species (37.5%) were exclusively found in the ‘northern/deep’, and 29 species (22.7%) were exclusively found in southern/shallow’ sampling sites (Figure 2a).

**Figure 2:**
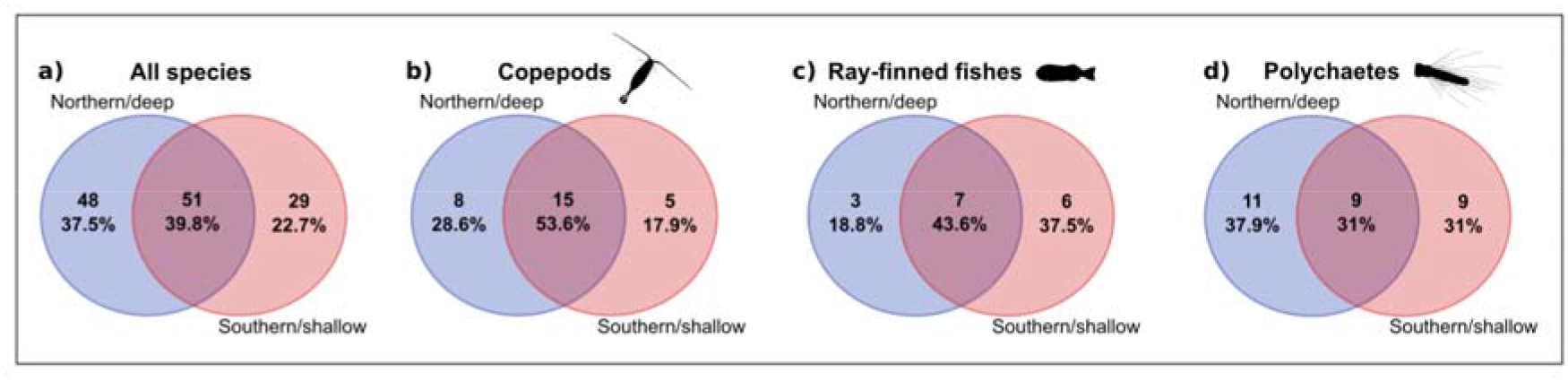
Number and percentage of identified species exclusively found in ‘northern/deep’ respectively ‘southern/shallow’ sampling sites, and number of species found in both areas. a) All species, b) Copepods, c) Ray-finned fishes, d) Polychaetes

### Copepods

We identified 28 copepod species in our dataset. Of these, 8 (28.6%) were found exclusively in the ‘northern/deep’ sites, 5 (17.9%) exclusively in the ‘southern/shallow’ sites, and 15 (53.6%) were found in both areas (Figure 2 b). The four most common species were *Oithona similis* (33.2% of all copepod reads), *Microcalanus pusillus* (13.3%), *Calanus finmarchicus* (12.6%), and *Temora longicornis* (11.4%).

We compared the metabarcoding data of these species to the abundance data recorded by the Continuous Plankton Recorder (CPR). For *Oithona similis*, analysis revealed no significant correlation of metabarcoding read abundance with latitude (Figure 3a), which is in congruence with the CPR data (Figure 3e). *Microcalanus pusillus* did not show a significant correlation of read abundance and latitude in the metabarcoding dataset (Figure 3b). However, high read abundances were found in central North Sea sampling sites (Site S109, 18.5% of copepod reads; Site S22, 45.4%), whereas abundance of the species did not exceed 6% of reads in all other sampling sites. Metabarcoding and CPR data could not be compared, as *Microcalanus pusillus* is absent from the CPR data due to its distribution in deeper water and small size (Fransz 1991). The CPR samples only the top water layer and the used mesh size does not reliably retain very small organisms. For *Calanus finmarchicus*, a strong, significant increase in read abundance with latitude was observed in the metabarcoding dataset (Figure 3c). Equally, the CPR dataset showed that abundance of the species significantly increased with latitude (Figure 3f). *Temora longicornis* showed a negative, but non-significant trend in read abundance from southern to northern sampling sites in the metabarcoding data (Figure 3d). This trend was also found in the CPR data and was significant (Figure 3g) (All statistical results: Tale S3)

**Figure 3:**
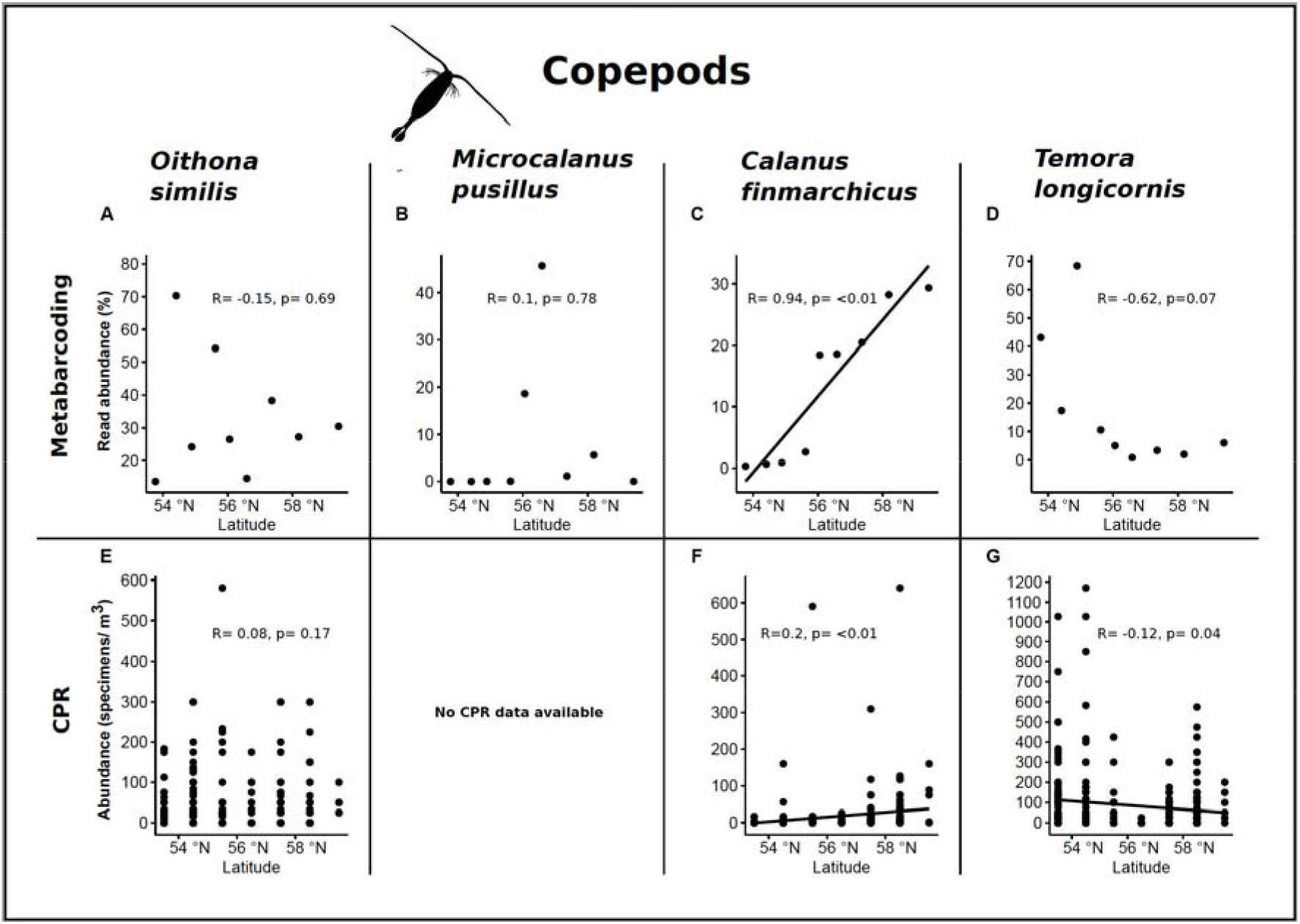
Correlations with latitude of the four most abundant copepod taxa in the metabarcoding dataset compared to data of the Continuous Plankton Recorder (CPR). Correlations were based on relative read abundances (metabarcoding data) and specimens/m^3^ (CPR).

### Ray-finned fishes

Out of 16 identified ray-finned fish species, 3 (18.8%) were found exclusively in the ‘northern/deep’ sampling sites, while 6 (37.5%) were only found in the ‘southern/shallow’ sites, and 7 (43.6%) were found in both areas (Figure 2c). The most abundant species were the common mackerel (*Scomber scombrus*, 48.6% of ray-finned fish reads), common dab (*Limanda limanda*, 19%), common ling (*Molva molva*, 11.8%) and the scaldfish *Arnoglossus laterna* (8.3%). As the CPR assesses fish on the level of eggs and larvae without species identification, no comparison of metabarcoding and CPR data was possible.

High read abundances of the common mackerel (*Scomber scombrus*) were found in the central and northern part of the North Sea (>90% of ray-finned fish reads; sites S109, 56.06 °N; S22, 56.59 °N; S93, 57.36 °N)(figure 4a). The common dab (*Limanda limanda*) was found in high abundance (>90% of fish reads) at sampling sites S176 (54.41°N) and S156 (54.89°N) in the southern part of the North Sea (figure 4b). For the common ling (*Molva molva*), a significant correlation of latitude and read abundance was found, with read abundance reaching >90% of reads in the northernmost sampling site S74 (59.42 °N, figure 4c). The scaldfish (*Arnoglossus laterna*) showed a high read abundance (>60%) in the southernmost sampling site S203 (53.77 °N), but no significant correlation of read abundance and latitude was found (figure 4d) (statistical results: table S4).

**Figure 4:**
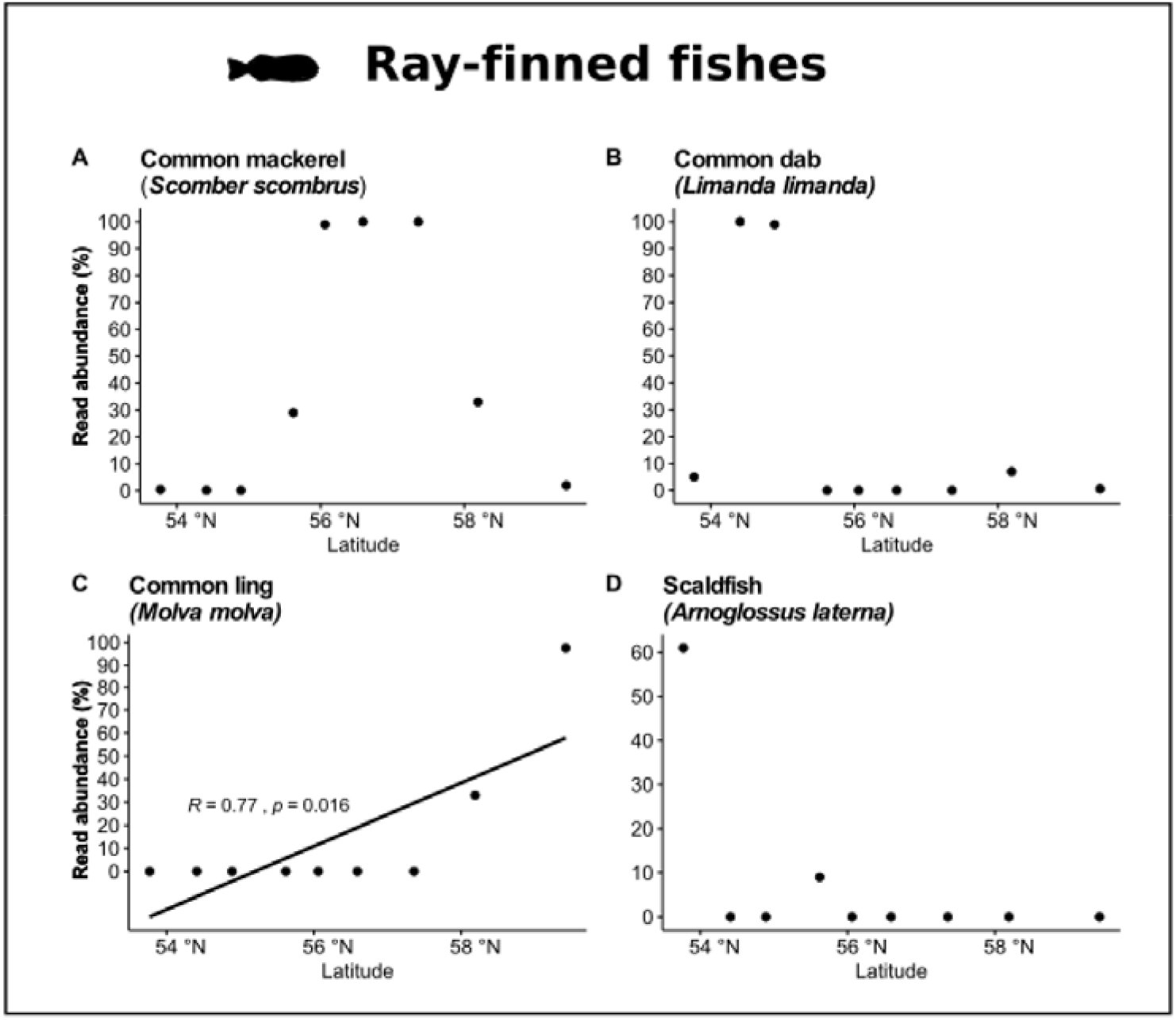
Correlations of relative read abundance with latitude of the four most abundant ray-finned fish species in the metabarcoding dataset (only shown when significant).

### Polychaetes

Of the 29 identified polychaete species, 11 (37.9%) were exclusively found in the ‘northern/deep’ sampling sites, 9 (31%) exclusively in the ‘southern/shallow’ sampling sites, and 9 (31%) were found in both areas (Figure 2 d). The most abundant species were *Paramphinome jeffreysii* (28.3% of polychaete reads), *Glycera lapidum* sp. 1 (20.9%), *Pectinaria koreni* (18.4%) and *Magelona johnstoni* (11.8%). As the CPR registers all polychaetes apart from the holoplanktonic Tomopteris, as (unidentified) larvae, no comparison of metabarcoding data and CPR data was possible.

*Paramphinome jeffreysii* showed a strong positive correlation of read abundance and latitude (figure 5 a), with read abundances reaching >60% of polychaete reads in all ‘northern/deep’ sampling sites, while it was only found in one site in the southern North Sea, with a read abundance of <1%. *Pectinaria koreni* did not show a significant correlation of read abundance and latitude, but was found in high abundance in two sampling sites in the ‘southern/shallow’ area of the North Sea (S156, 47%; S130, 29%), and 20% read abundance in the northernmost sampling site (figure 5b). *Glycera lapidum* sp. 1 was commonly found with less than 1% of reads, except in the two southern sampling sites S176 (54.41°N) and S130 (55.62°N), where the species occurred with >50% read abundance. No significant correlation of read abundance and latitude was found (figure 5c). For *Magelona johnstoni*, the highest relative abundances were found in ‘southern/shallow’ sites (S203, 63%; respectively 32% in S156). Read abundance at all other sampling sites was <1% (figure 5d) (All statistical results: Table S5)

**Figure 5:**
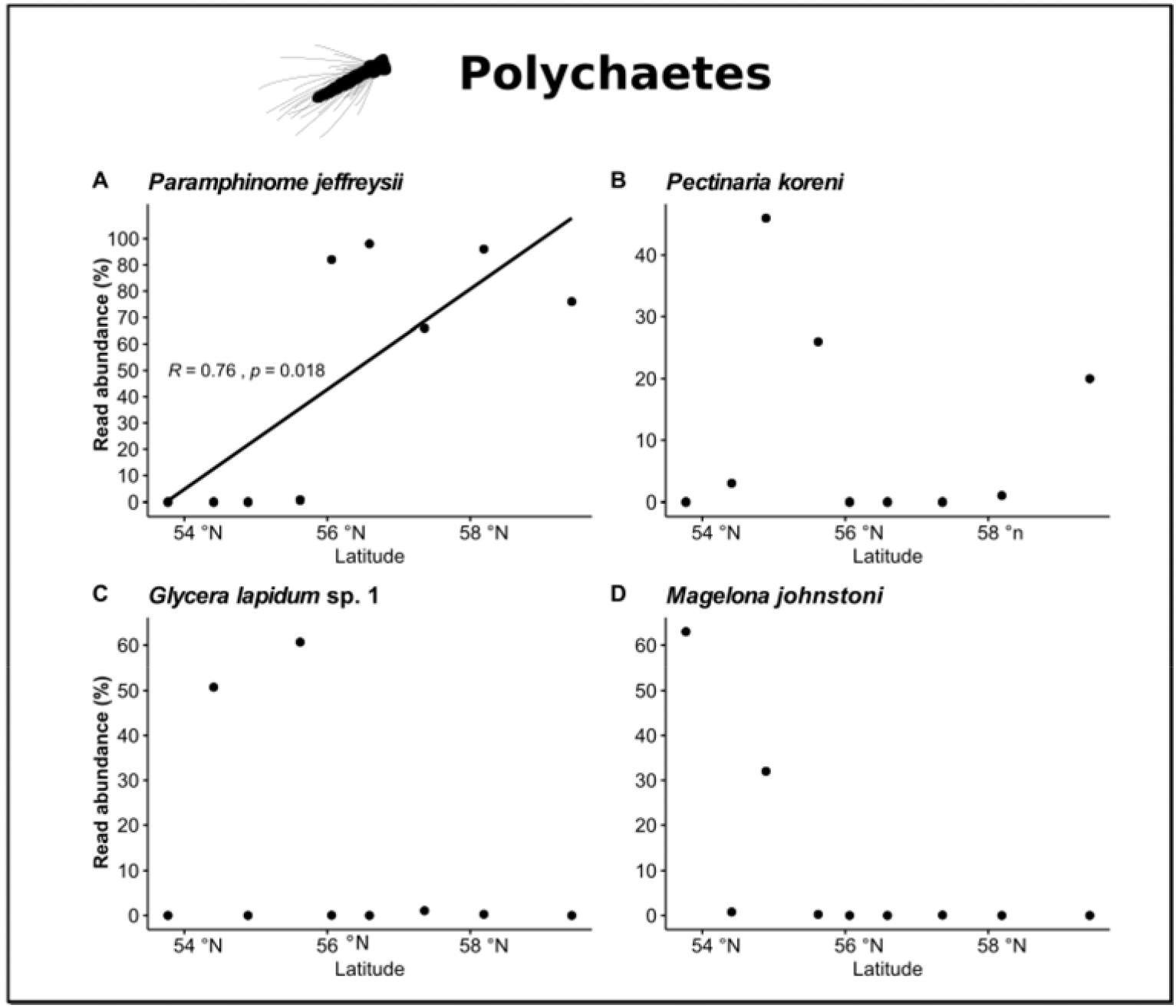
Correlations of relative read abundance with latitude of the four most abundant polychaete species in the metabarcoding dataset (only shown when significant).

## Discussion

Using metabarcoding we show that the overall community composition of zooplankton differs between northern and southern areas of the North Sea. Differences in community composition are associated with the influx of Atlantic water from the north, which has been documented before based on morphological identifications of zooplankton (Fransz et al., 1991; Krause, M., Dippner, J.W., Beil, J., 1995). However, the eggs and larval stages of common groups such as fish and polychaetes are difficult to identify based on morphology, which is why they are often not recorded by long-term monitoring programs such as the CPR (Reid et al., 2003a). Metabarcoding can help identify those species, potentially allowing better insights into the composition of communities and species distribution patterns (Kimmerling et al., 2018; Schroeder et al., 2020). Our data show that valuable information on the distribution of zooplankton in the North Sea can be gained through metabarcoding, including information of meroplanktonic species that are still little studied. However, we acknowledge that our study is based on relatively few samping sites. To assess the distribution of zooplankton species in the North Sea more accurately a denser sampling scheme in time and space should be employed. Such sampling is performed by the Continuous Plankton Recorder (Reid et al., 2003a), and future studies should aim at utilising CPR samples for metabarcoding (see e.g. (Kirby, Lindley & Batten, 2006; Stern et al., 2018)). In this way, the long-term CPR dataset could also provide more detailed insights into meroplanktonic groups such as fish and benthic invertebrates. In this study, we could only compare our metabarcoding results to the CPR data on copepods. We point out that comparisons of metabarcoding and CPR data should be interpreted with care, as the CPR samples only the top layers of the water column, and abundance inferred from metabarcoding data can be biased due to amplification and primers bias (Elbrecht & Leese, 2015, 2017; Wangensteen et al., 2018; Zizka et al., 2019a). Further, comparing metabarcoding read abundance and number of specimens, as mostly provided by previous morphological studies, is challenging, as reads do not automatically correlate with specimen numbers. However, our results show that the abundance data inferred through metabarcoding mostly matches with the known distribution of species described in previous studies, thus underlining that metabarcoding is a potentially powerful tool for monitoring of zooplankton communities. The highly degenerate Leray XT primers used in this study amplified a wide range of planktonic taxa. A high percentage (96%) of reads could be assigned to a taxonomic class, and only 1% of these reads were assigned to non-metazoan taxa. This is in contrast to previous metabarcoding studies targeting highly diverse communities, which highlighted that degenerate primers commonly co-amplify a high number of non-metazoan taxa (Weigand & Macher, 2018; Wangensteen et al., 2018). We assume that only little non-metazoan biomass was present in our samples, leading to successful amplification of the target planktonic animals. However, only 75% of reads could be assigned to species level, which highlights the lack of genetic reference data for marine planktonic organisms. This can render molecular identification of species impossible (see e.g. (Porter & Hajibabaei, 2018; Weigand et al., 2019)) and underlines the need for improved reference databases, which can only be otained with the help of taxonomic experts. We discuss the findings on copepods, ray-finned fishes and polychaetes in more detail below.

### Copepods

Our results show that the copepod community significantly differs between northern and southern areas of the North Sea, which is congruent with previous studies based on morphological identifications (Fransz et al., 1991; Beaugrand et al., 2002). Comparison of the most abundant copepod species in our metabarcoding dataset with CPR data showed a high level of congruence. *Oithona similis* did not show significant differences in read abundance between ‘northern/deep’ and ‘southern/shallow’ sampling sites, which corresponds to the *Oithona* spp. abundances recorded by the CPR. We acknowledge that a direct comparison of Oithona similis and *Oithona* spp. data are potentially biased, as other *Oithona* species can be present in the North Sea. Fransz (1991) however pointed out that *O. similis* is commonly the most abundant *Oithona* species in the North Sea. Corresponding to that information, the only other *Oithona* species we identified in our metabarcoding dataset was *Oithona atlantica*, which was found in low read abundances (1% of copepod reads). We therefore see the comparison as legitimate. *Microcalanus pusillus* was the second most abundant species in the metabarcoding dataset, but we could not compare our data with CPR data. The species is likely overlooked by CPR sampling due to its distribution in deeper water layers and its small size, which are not sampled by the CPR (Hays & Warner, 1993; Hays, 1994). Previous work identified *M. pusillus* as a mostly Atlantic species (Fransz et al., 1991; Beaugrand et al., 2002) but our data did not show a correlation between abundance of this species and latitude. We found *M. pusillus* in high abundance in the central North Sea. This area, which is located north of the shallow Doggers Bank is known for upwellings bringing nutrient-rich bottom water closer to the surface (Nielsen et al., 1993), which might explain the high abundance of this species in the area. *Calanus finmarchicus* showed a strong positive correlation of abundance with latitude, and this pattern was also found in the CPR data. Our results are in congruence with previous findings, which found the species to be mostly restricted to northern, Atlantic waters (Beaugrand et al., 2002; Marshall & Orr, 2013). For *Temora longicornis*, an indicator species of coastal waters (Beaugrand et al., 2002), high abundances in the coastal regions were found in both the metabarcoding dataset and the CPR data.

Copepod species that we found exclusively in the ‘northern/deep’ area of the North Sea, such as *Candacia armata*, *Scolecithricella minor*, *Anomalocera patersoni*, *Diaixis hibernica* and *Pseudocalanus acuspes*, are all species known predominantly from northern areas of the North Sea (Fransz et al., 1991; Beaugrand et al., 2002; Hovda & Fosshagen, 2003). We further identified *Goniopsyllus rostrata* and *Clausocalanus pergens*, which to the best of our knowledge have not yet been reported from the North Sea. The reference sequences in the BOLD database stem from specimens sampled off northern Spain (BOLD BIN numbers: AAO2968; AAJ1005). We further detected *Pseudocalanus mimus*, which is generally considered a North Pacific species (Frost, 1989; Questel et al., 2016), and that is also where the BOLD references (BIN AAH8134) stem from. However, a few records of the species from between Canada and Greenland exist (http://www.iobis.org/)(Nelson, 2014). The species has not previously been recorded from the North East Atlantic, which could mean that the species was either overlooked, or that its distribution range has recently expanded. Of the copepod species that were exclusively found in the southern sampling sites, all except *Caligus elongatus*, a fish parasite, are known as typical of coastal/shallow waters: *Longipedia* sp. DZMB181 (Khodami et al., 2017), *Haloschizopera pygmaea* (Rossel & Martínez Arbizu, 2019)*, Isias clavipes (Beaugrand et al., 2000*), and *Acartia tonsa* (Fransz et al., 1991; Caudill & Bucklin, 2004).

### Ray-finned fishes

Fish larvae and eggs are part of the zooplankton for a limited time and their occurrence is mostly influenced by spawning and nursery areas to which eggs and larvae drift (Knijn, 1993; Gibson, 2001; Gibson et al., 2015). We found that the inferred community composition of ray-finned fishes strongly differed between northern and southern sampling sites in the North Sea, and that our data corresponds to known distribution patterns of fish species and their spawning areas. The three species exclusively found in the ‘northern/deep’ sampling sites were the grey gurnard (*Eutrigla gurnadus*), the argentine (*Argentina sphyraena*) and the slender snipe eel (*Nemichthys scolopaceus*). The grey gurnard and the argentine are known to occur mostly in deeper, northern waters (Knijn, 1993; Wright, Jensen & Tuck, 2000), while the slender snipe eel is known from deep sea environments (Feagans-Bartow & Sutton, 2014; Lusher et al., 2016). This corresponds to our results, as we found the slender snipe eel exclusively in the Devil’s Hole sampling site (S22) which reaches 230m water depth. Of the species exclusively found in ‘southern/shallow’ sampling sites, the solenette (*Buglossidium luteum*), European sprat (*Sprattus sprattus*), striped red mullet (*Mullus surmuletus*) and common sole (*Solea solea*) are all known to be mainly distributed along the coastlines and in southern regions of the North Sea (Knijn, 1993; Milner, 2016), which is congruent with our findings. The only exception is the sand eel *Ammodytes marinus*, which can be found in shallow, sandy habitats throughout the North Sea, but we found in only one sampling site in the southern North Sea. We assume that we did not find the species in more samples due to low overall abundance and competition with other species during amplification and sequencing of the data. Separate analyses of the four most abundant ray-finned fish taxa correspond to previous findings showing that coastal areas of the North Sea are important spawning and nursing grounds for these species, namely, common dab (*Limanda limanda*) (Bolle et al., 1994) and scaldfish (*Arnoglossus laterna*)(Land & Van der Land, 1991; van Hal, Smits & Rijnsdorp, 2010). Our finding that the common mackerel (*Scomber scombrus*) showed the highest read abundance in the central part of the North Sea corresponds to the known spawning area of this species (Jansen et al., 2012). The common ling (*Molva molva*), found in high abundances in the northernmost sampling sites, is also known to spawn in these areas (Knijn, 1993). Further research will show if metabarcoding will detect known distribution and spawning areas for a high number of fish species, which will be helpful for monitoring of populations.

### Polychaetes

With the exception of the holoplanktonic *Tomopteris spp.*, polychaetes in the North Sea are meroplanktonic (Plate & Husemann, 1994; Van Ginderdeuren et al., 2014). Even though polychaetes are a highly diverse and abundant group, their planktonic stages are relatively little known due to difficulties in identification (Williams et al., 1993; Vezzulli & Reid, 2003; Heimeier, Lavery & Sewell, 2010). As for the copepods and ray-finned fishes, we found a strong difference in community composition of polychaetes between northern and southern sampling sites in the North Sea. This corresponds to known patterns of macrobenthos community differences between shallow areas in the southern and northern North Sea (Duineveld et al., 1991). However, information on the distribution of most of the identified species in the North Sea is scarce or not available, rendering a comparison of our metabarcoding data to previous data based on morphological identifications mostly impossible. We assume that the lack of information on many species is due to difficulties in reliable identification and the lack of taxonomic experts, which highlights the need for a combined morphological and molecular approach for future studies and the preparation of reference libraries. Separate analyses of the four most abundant species showed that the polychaete community was dominated by *Paramphinome jeffreysii* in the ‘northern/deep’ sampling sites. In congruence with our results, this species was previously found in high abundance in northern regions (Kröncke et al., 2011). *Pectinaria koreni* has been recorded in areas of fine sediment, often closer to the coast (Thiébaut et al., 1997; Desroy, 2003) and from areas near the Shetland islands (GBIF dataset: https://doi.org/10.15468/39omei). We also found this species in high abundance in the ‘southern/shallow’ area of the North Sea, as well as in the northernmost sampling site close to the Shetland Islands. Little information is available on the distribution and larval stages of *Glycera lapidum* and *Magelona johnstoni*. Both species are known from several regions of the North Sea (Kunitzer et al., 1992; Meißner & Darr, 2009). We consider it possible that the high abundance of these species in a few sampling sites can be explained by local spawning events. Overall, our results show the power of metabarcoding to assess the meroplanktonic polychaete community, but we conclude that more combined molecular and morphological work is required to fully understand distribution patterns of polychaete larvae.

### Conclusion

We showed that metabarcoding of zooplankton samples from the North Sea, using highly degenerate COI primers, can give valuable insights into the diversity and distribution of planktonic animals. We found clear differences in the overall zooplankton assemblages between northern and southern areas of the North Sea, as well as more specifically for copepods, ray-finned fishes and polychaetes. Our results were largely congruent with previous studies based on morphological identifications, which indicates the robustness of our molecular approach. Nevertheless, we highlight the need for more complete reference databases to be able to make full use of the information gained through metabarcoding. We suggest that metabarcoding should be considered for implementation into future biodiversity assessments, as the ability to quickly assess whole zooplankton samples is valuable for biodiversity studies in times of rapid ocean changes.

## Acknowledgements

We thank the Pelagia crew for help during the NICO leg 10 cruise and Rob Witbaard for organising and leading the NICO leg 10. We thank Elza Duijm for support in the lab.

## Data availability

All raw data are available from figshare: https://doi.org/10.6084/m9.figshare.12698054.v1

## Field Study Permissions

The following information was supplied relating to field study approvals (i.e., approving body and any reference numbers): Sampling in UK waters was approved by the Maritime Policy Unit (Legal Directorate) of the Foreign and Commonwealth Office (ref 33/2018).

