## Supplementary tables for "Metabarcoding reveals different zooplankton communities in northern and southern areas of the North Sea"

**Supplementary tables 1-5**

**Table S1: Sampling stations and recorded abiotic variables recorded during the NICO 10 expedition from the Dutch Coast to the Shetland Islands**

| Sampling site name | Coordinates (°N, °E) | Mean remperature (°C) | Mean salinity (PSU) | Depth (m) |
|---|---|---|---|---|
| **S74** | 59.416510, 0.499900 | 8.2 | 35.1 | 134 |
| **S37** | 58.1855556, 0.5016667 | 8.7 | 35.1 | 89 |
| **S93** | 57.36046, 0.57784 | 7.8 | 34.8 | 84 |
| **S22** | 56.5866667, 0.6905556 | 8.3 | 34.9 | 220 |
| **S109** | 56.06489, 1.59652 | 8.7 | 35 | 79 |
| **S130** | 55.62157, 2.38651 | 7.8 | 34.8 | 73 |
| **S156** | 54.88581, 3.69192 | 8.3 | 34.6 | 41 |
| **S176** | 54.41489, 4.04154 | 9.6 | 34.6 | 43 |
| **S203** | 53.76851, 4.76715 | 11.8 | 34.5 | 34 |

**Table S2: Species list and read number per sampling site**

| Class | Order | Family | Genus | Species | S22 | S37 | S74 | S93 | S109 | S130 | S156 | S176 | S203 |
|---|---|---|---|---|---|---|---|---|---|---|---|---|---|
| Copepoda | Calanoida | Acartiidae | Acartia | Acartia clausi | 0 | 0 | 0 | 72 | 0 | 170 | 15 | 630 | 3995 |
| Copepoda | Calanoida | Acartiidae | Acartia | Acartia tonsa | 0 | 0 | 0 | 0 | 0 | 0 | 0 | 0 | 23 |
| Hydrozoa | Trachymedusae | Rhopalonematidae | Aglantha | Aglantha digitale | 0 | 0 | 0 | 0 | 1870 | 117 | 420 | 629 | 0 |
| Actinopterygii | Trachiniformes | Ammodytidae | Ammodytes | Ammodytes marinus | 0 | 0 | 0 | 0 | 0 | 263 | 0 | 35 | 0 |
| Copepoda | Harpacticoida | Miraciidae | Amphiascopsis | Amphiascopsis cinct | 344 | 0 | 0 | 992 | 2477 | 2500 | 9574 | 8947 | 0 |
| Ophiuroidea | Amphilepidida | Amphiuridae | Amphiura | Amphiura filiformis | 0 | 0 | 0 | 0 | 219 | 0 | 0 | 1470 | 63233 |
| Copepoda | Calanoida | Pontellidae | Anomalocera | Anomalocera paterso | 0 | 0 | 586 | 0 | 0 | 0 | 0 | 0 | 0 |
| Bivalvia | Venerida | Arcticidae | Arctica | Arctica islandica | 0 | 0 | 29 | 0 | 0 | 0 | 0 | 0 | 0 |
| Actinopterygii | Argentiniformes | Argentinidae | Argentina | Argentina sphyraena | 0 | 841 | 0 | 0 | 0 | 0 | 0 | 0 | 0 |
| Actinopterygii | Pleuronectiformes | Bothidae | Arnoglossus | Arnoglossus laterna | 0 | 40 | 0 | 0 | 0 | 37 | 0 | 0 | 389949 |
| Asteroidea | Forcipulatida | Asteriidae | Asterias | Asterias rubens | 0 | 0 | 368 | 0 | 0 | 0 | 77 | 528 | 425 |
| Asteroidea | Paxillosida | Astropectinidae | Astropecten | Astropecten irregular | 0 | 0 | 46 | 0 | 0 | 0 | 0 | 0 | 0 |
| Asteroidea | Paxillosida | Astropectinidae | Astropecten | Astropecten irregular | 0 | 0 | 0 | 0 | 119 | 0 | 0 | 440 | 412 |
| Thecostraca | Sessilia | Balanidae | Balanus | Balanus balanus | 0 | 0 | 0 | 3655 | 2807 | 2851 | 0 | 0 | 0 |
| Thecostraca | Sessilia | Balanidae | Balanus | Balanus crenatus | 82 | 0 | 0 | 0 | 0 | 0 | 0 | 0 | 0 |
| Leptocardii | Amphioxiformes | Branchiostomatidae | Branchiostoma | Branchiostoma lance | 0 | 0 | 0 | 0 | 38 | 0 | 0 | 0 | 0 |
| Echinoidea | Spatangoida | Brissidae | Brissopsis | Brissopsis lyrifera | 0 | 528 | 36554 | 107 | 0 | 413 | 0 | 0 | 0 |
| Actinopterygii | Pleuronectiformes | Soleidae | Buglossidium | Buglossidium luteum | 0 | 0 | 0 | 0 | 0 | 0 | 0 | 0 | 207835 |
| Copepoda | Calanoida | Calanidae | Calanus | Calanus euxinus | 12831 | 881 | 2158 | 5604 | 10297 | 0 | 40 | 188 | 471 |
| Copepoda | Calanoida | Calanidae | Calanus | Calanus finmarchicus | 218066 | 79775 | 100711 | 83487 | 156808 | 37719 | 2198 | 1824 | 1341 |
| Copepoda | Calanoida | Calanidae | Calanus | Calanus helgolandicu | 136255 | 10209 | 20736 | 38396 | 156915 | 1273 | 833 | 8596 | 4763 |
| Copepoda | Siphonostomatoida | Caligidae | Caligus | Caligus elongatus | 0 | 0 | 0 | 0 | 0 | 0 | 0 | 127 | 0 |
| Actinopterygii | Callionymiformes | Callionymidae | Callionymus | Callionymus reticulat | 0 | 0 | 0 | 0 | 0 | 0 | 0 | 0 | 64 |
| Copepoda | Calanoida | Candaciidae | Candacia | Candacia armata | 2085 | 28786 | 2155 | 5691 | 1102 | 0 | 0 | 0 | 0 |
| Copepoda | Calanoida | Centropagidae | Centropages | Centropages hamatu | 0 | 0 | 372 | 32 | 0 | 0 | 0 | 0 | 403 |
| Copepoda | Calanoida | Centropagidae | Centropages | Centropages typicus | 387 | 67 | 820 | 122 | 90 | 8 | 0 | 0 | 335 |
| Polychaeta | Phyllodocida | Nereididae | Ceratocephale | Ceratocephale loveni | 0 | 0 | 0 | 0 | 0 | 19 | 0 | 0 | 0 |
| Pilidiophora | Heteronemertea | Lineidae | Cerebratulus | Cerebratulus sp. Cm | 0 | 1942 | 284 | 0 | 0 | 0 | 0 | 0 | 0 |
| Polychaeta | Canalipalpata | Chaetopteridae | Chaetopterus | Chaetopterus sarsi | 0 | 0 | 47 | 0 | 0 | 47 | 34470 | 15008 | 40 |
| Copepoda | Calanoida | Clausocalanidae | Clausocalanus | Clausocalanus perge | 0 | 0 | 0 | 0 | 41 | 0 | 0 | 0 | 0 |
| Gastropoda | Pteropoda | Clionidae | Clione | Clione limacina | 0 | 0 | 0 | 0 | 30 | 273 | 42 | 0 | 0 |
| Malacostraca | Decapoda | Corystidae | Corystes | Corystes cassivelau | 0 | 0 | 0 | 0 | 35 | 0 | 361 | 2345 | 983 |
| Scyphozoa | Semaeostomeae | Cyaneidae | Cyanea | Cyanea capillata | 0 | 0 | 0 | 0 | 26 | 0 | 0 | 0 | 0 |
| Gastropoda | Cephalaspidea | Cylichnidae | Cylichna | Cylichna cylindracea | 0 | 0 | 0 | 0 | 0 | 0 | 0 | 0 | 162 |
| Copepoda | Calanoida | Diaixidae | Diaixis | Diaixis hibernica | 623 | 0 | 0 | 0 | 0 | 0 | 0 | 0 | 0 |
| Echinoidea | Spatangoida | Loveniidae | Echinocardium | Echinocardium corda | 271460 | 196 | 1231 | 8758 | 24231 | 750 | 2217 | 246711 | 363200 |
| Echinoidea | Spatangoida | Loveniidae | Echinocardium | Echinocardium flaves | 2192 | 0 | 3372 | 1502 | 29079 | 418 | 494 | 1136 | 0 |
| Gastropoda | Nudibranchia | Eubranchidae | Eubranchus | Eubranchus exiguus | 0 | 0 | 0 | 0 | 45 | 44 | 0 | 0 | 0 |
| Sagittoidea | Phragmophora | Eukrohniidae | Eukrohnia | Eukrohnia hamata | 1750 | 6824 | 90 | 1429 | 189 | 34 | 826 | 0 | 139 |
| Polychaeta | Phyllodocida | Phyllodocidae | Eumida | Eumida bahusiensis | 29 | 0 | 0 | 0 | 0 | 0 | 0 | 0 | 0 |
| Polychaeta | Phyllodocida | Nereididae | Eunereis | Eunereis longissima | 0 | 0 | 0 | 0 | 0 | 0 | 0 | 239 | 18772 |
| Actinopterygii | Scorpaeniformes | Triglidae | Eutrigla | Eutrigla gurnardus | 2760 | 0 | 0 | 0 | 3159 | 0 | 0 | 0 | 0 |
| Branchiopoda | Onychopoda | Podonidae | Evadne | Evadne nordmanni | 82 | 880249 | 718586 | 596 | 10678 | 112196 | 56 | 1594 | 81925 |
| Bivalvia | Cardiida | Tellinidae | Fabulina | Fabulina fabula | 0 | 0 | 0 | 0 | 142 | 65 | 0 | 0 | 160 |
| Polychaeta | Phyllodocida | Polynoidae | Gattyana | Gattyana cirrhosa | 0 | 0 | 0 | 0 | 0 | 0 | 0 | 109 | 0 |
| Enteropneusta | Enteropneusta_order_incertae_sedis | Ptychoderidae | Glossobalanus | Glossobalanus marg | 0 | 0 | 4786 | 0 | 7676 | 0 | 0 | 0 | 0 |
| Polychaeta | Phyllodocida | Glyceridae | Glycera | Glycera lapidum I | 0 | 26 | 0 | 169 | 31 | 98007 | 0 | 114482 | 0 |
| Polychaeta | Phyllodocida | Glyceridae | Glycera | Glycera lapidum II | 0 | 130 | 0 | 0 | 0 | 0 | 0 | 0 | 0 |
| Actinopterygii | Pleuronectiformes | Pleuronectidae | Glyptocephalus | Glyptocephalus cyno | 0 | 0 | 38 | 0 | 0 | 0 | 0 | 1277 | 0 |
| Copepoda | Harpacticoida | Peltidiidae | Goniopsyllus | Goniopsyllus rostrata | 0 | 0 | 0 | 33 | 0 | 0 | 0 | 0 | 0 |
| Echinoidea | Camarodonta | Echinidae | Gracilechinus | Gracilechinus acutus | 0 | 75 | 12007 | 36 | 47 | 59 | 0 | 0 | 0 |
| Copepoda | Harpacticoida | Miraciidae | Haloschizopera | Haloschizopera pygr | 0 | 0 | 0 | 0 | 0 | 0 | 0 | 161 | 0 |
| Polychaeta | Phyllodocida | Polynoidae | Harmothoe | Harmothoe glabra | 0 | 0 | 0 | 0 | 146 | 0 | 0 | 0 | 0 |
| Actinopterygii | Pleuronectiformes | Pleuronectidae | Hippoglossoides | Hippoglossoides plat | 0 | 361994 | 1131 | 0 | 85 | 0 | 0 | 82 | 0 |
| Malacostraca | Decapoda | Oregoniidae | Hyas | Hyas coarctatus | 1929 | 0 | 0 | 4590 | 0 | 0 | 0 | 0 | 0 |
| Copepoda | Calanoida | Centropagidae | Isias | Isias clavipes | 0 | 0 | 0 | 0 | 0 | 62 | 0 | 0 | 0 |
| Pilidiophora | Heteronemertea | Cerebratulidae | Leucocephalonemertes | Leucocephalonemert | 0 | 0 | 4793 | 0 | 126 | 0 | 0 | 0 | 0 |
| Actinopterygii | Pleuronectiformes | Pleuronectidae | Limanda | Limanda limanda | 110 | 93018 | 701 | 69 | 33 | 0 | 236360 | 527889 | 34446 |
| Malacostraca | Decapoda | Polybiidae | Liocarcinus | Liocarcinus holsatus | 0 | 0 | 0 | 0 | 0 | 0 | 0 | 0 | 5218 |
| Hydrozoa | Anthoathecata | Rathkeidae | Lizzia | Lizzia blondina | 0 | 0 | 655 | 0 | 0 | 0 | 0 | 0 | 0 |
| Copepoda | Canuelloida | Longipediidae | Longipedia | Longipedia coronata | 0 | 0 | 238 | 39 | 0 | 410 | 614 | 7334 | 0 |
| Copepoda | Canuelloida | Longipediidae | Longipedia | Longipedia sp. DZME | 0 | 0 | 0 | 0 | 0 | 701 | 2332 | 780 | 1283 |
| Asteroidea | Paxillosida | Luidiidae | Luidia | Luidia sarsi | 0 | 0 | 165 | 0 | 170 | 84 | 0 | 0 | 0 |
| Bivalvia | Venerida | Mactridae | Mactra | Mactra stultorum | 0 | 0 | 0 | 0 | 0 | 0 | 0 | 53 | 0 |
| Polychaeta | Spionida | Magelonidae | Magelona | Magelona johnstoni | 0 | 0 | 0 | 14 | 0 | 421 | 81193 | 1730 | 37048 |
| Polychaeta | Spionida | Magelonidae | Magelona | Magelona mirabilis | 0 | 0 | 0 | 0 | 0 | 0 | 736 | 28 | 134 |
| Malacostraca | Euphausiacea | Euphausiidae | Meganyctiphanes | Meganyctiphanes no | 102 | 0 | 0 | 85 | 0 | 0 | 0 | 0 | 0 |
| Hydrozoa | Leptothecata | Melicertidae | Melicertum | Melicertum octocosta | 0 | 0 | 0 | 0 | 0 | 0 | 40 | 0 | 0 |
| Gymnolaemata | Cheilostomatida | Membraniporidae | Membranipora | Membranipora memb | 0 | 0 | 46 | 0 | 0 | 0 | 0 | 0 | 0 |
| Actinopterygii | Gadiformes | Gadidae | Merlangius | Merlangius merlangu | 87 | 0 | 39 | 0 | 0 | 0 | 1609 | 0 | 0 |
| Copepoda | Calanoida | Metridinidae | Metridia | Metridia lucens | 810 | 15 | 1705 | 161 | 1738 | 0 | 30 | 0 | 0 |
| Copepoda | Calanoida | Clausocalanidae | Microcalanus | Microcalanus pusillus | 536595 | 16008 | 88 | 4638 | 158236 | 604 | 72 | 0 | 0 |
| Actinopterygii | Gadiformes | Gadidae | Molva | Molva molva | 0 | 438332 | 116697 | 0 | 75 | 0 | 0 | 0 | 29 |
| Actinopterygii | Perciformes | Mullidae | Mullus | Mullus surmuletus | 0 | 0 | 0 | 0 | 0 | 0 | 0 | 0 | 322 |
| Actinopterygii | Anguilliformes | Nemichthyidae | Nemichthys | Nemichthys scolopac | 57 | 0 | 0 | 0 | 0 | 0 | 0 | 0 | 0 |
| Malacostraca | Decapoda | Nephropidae | Nephrops | Nephrops norvegicus | 0 | 0 | 0 | 313 | 0 | 0 | 0 | 0 | 0 |
| Polychaeta | Phyllodocida | Hesionidae | Nereimyra | Nereimyra punctata | 0 | 0 | 0 | 1375 | 0 | 0 | 0 | 0 | 0 |
| Polychaeta | Phyllodocida | Nereididae | Nereis | Nereis elitoralis | 0 | 0 | 0 | 0 | 0 | 0 | 17 | 253 | 1104 |
| Copepoda | Cyclopoida | Oithonidae | Oithona | Oithona atlantica | 12740 | 150 | 5904 | 5181 | 24630 | 4099 | 345 | 191 | 0 |
| Copepoda | Cyclopoida | Oithonidae | Oithona | Oithona similis | 150397 | 77623 | 99917 | 156311 | 214431 | 805053 | 54499 | 188175 | 52589 |
| Gastropoda | Nudibranchia | Onchidorididae | Onchidoris | Onchidoris muricata | 0 | 0 | 0 | 57 | 0 | 0 | 0 | 0 | 0 |
| Gastropoda | Littorinimorpha | Rissoidae | Onoba | Onoba semicostata | 0 | 0 | 0 | 1735 | 57 | 0 | 0 | 0 | 0 |
| Ophiuroidea | Amphilepidida | Ophiotrichidae | Ophiothrix | Ophiothrix fragilis | 0 | 0 | 0 | 0 | 0 | 0 | 0 | 0 | 32 |
| Polychaeta | Eunicida | Dorvilleidae | Ophryotrocha | Ophryotrocha macula | 0 | 0 | 0 | 0 | 833 | 0 | 0 | 0 | 0 |
| Polychaeta | Sabellida | Oweniidae | Owenia | Owenia fusiformis | 0 | 0 | 185 | 0 | 0 | 0 | 0 | 0 | 0 |
| Polychaeta | Phyllodocida | Hesionidae | Oxydromus | Oxydromus vittatus | 0 | 0 | 3361 | 0 | 0 | 0 | 0 | 0 | 0 |
| Pilidiophora | Heteronemertea | Baseodiscidae | Oxypolia | Oxypolia beaumontia | 217 | 21404 | 30 | 2287 | 0 | 0 | 0 | 0 | 0 |
| Malacostraca | Decapoda | Paguridae | Pagurus | Pagurus bernhardus | 0 | 0 | 0 | 0 | 43 | 0 | 0 | 0 | 0 |
| Copepoda | Calanoida | Paracalanidae | Paracalanus | Paracalanus parvus | 38306 | 3207 | 5537 | 4654 | 42423 | 91425 | 1156 | 172 | 156246 |
| Polychaeta | Amphinomida | Amphinomidae | Paramphinome | Paramphinome jeffre | 115633 | 10936 | 67677 | 12403 | 79533 | 1369 | 0 | 0 | 0 |
| Polychaeta | Terebellida | Pectinariidae | Pectinaria | Pectinaria auricoma | 0 | 0 | 71 | 0 | 0 | 0 | 0 | 0 | 0 |
| Polychaeta | Terebellida | Pectinariidae | Pectinaria | Pectinaria koreni | 0 | 165 | 17591 | 0 | 0 | 47076 | 116659 | 5886 | 0 |
| Polychaeta | Phyllodocida | Pholoidae | Pholoe | Pholoe baltica | 0 | 0 | 0 | 0 | 0 | 0 | 16386 | 77182 | 0 |
| Phoronida_class_incertae_sedis | Phoronida_order_incertae_sedis | Phoronidae | Phoronis | Phoronis muelleri | 0 | 0 | 3989 | 0 | 525 | 0 | 0 | 0 | 0 |
| Polychaeta | Phyllodocida | Phyllodocidae | Phyllodoce | Phyllodoce groenland | 0 | 0 | 0 | 0 | 0 | 38 | 0 | 0 | 0 |
| Polychaeta | Terebellida | Terebellidae | Pista | Pista cristata | 452 | 0 | 0 | 0 | 149 | 978 | 952 | 95 | 291 |
| Gastropoda | unranked | Limapontiidae | Placida | Placida dendritica | 0 | 0 | 0 | 0 | 0 | 19 | 0 | 0 | 0 |
| Branchiopoda | Onychopoda | Podonidae | Pleopis | Pleopis polyphemoide | 0 | 0 | 0 | 0 | 0 | 9494 | 44 | 0 | 4309 |
| Branchiopoda | Onychopoda | Podonidae | Podon | Podon intermedius | 0 | 0 | 136 | 0 | 0 | 0 | 0 | 0 | 0 |
| Polychaeta | Spionida | Poecilochaetidae | Poecilochaetus | Poecilochaetus serpe | 0 | 0 | 0 | 0 | 0 | 0 | 0 | 0 | 1438 |
| Malacostraca | Decapoda | Polybiidae | Polybius | Polybius henslowii | 318 | 0 | 0 | 0 | 0 | 0 | 0 | 0 | 2372 |
| Polychaeta | Canalipalpata | Polygordiidae | Polygordius | Polygordius appendic | 0 | 0 | 660 | 0 | 0 | 0 | 0 | 0 | 0 |
| Malacostraca | Decapoda | Crangonidae | Pontophilus | Pontophilus spinosus | 0 | 0 | 0 | 0 | 197 | 0 | 0 | 0 | 0 |
| Asteroidea | Valvatida | Poraniidae | Porania | Porania pulvillus | 0 | 0 | 973 | 0 | 0 | 0 | 0 | 0 | 0 |
| Polychaeta | Spionida | Spionidae | Prionospio | Prionospio cirrifera | 0 | 128 | 0 | 0 | 0 | 0 | 0 | 0 | 0 |
| Malacostraca | Decapoda | Processidae | Processa | Processa modica | 0 | 855 | 0 | 0 | 0 | 0 | 0 | 0 | 0 |
| Copepoda | Calanoida | Clausocalanidae | Pseudocalanus | Pseudocalanus acus | 23 | 0 | 0 | 0 | 0 | 0 | 0 | 0 | 0 |
| Copepoda | Calanoida | Clausocalanidae | Pseudocalanus | Pseudocalanus mimu | 35 | 0 | 0 | 0 | 0 | 0 | 0 | 0 | 0 |
| Copepoda | Calanoida | Clausocalanidae | Pseudocalanus | Pseudocalanus moul | 1637 | 32 | 714 | 73 | 4069 | 1301 | 29 | 3580 | 2154 |
| Polychaeta | Sabellida | Sabellariidae | Sabellaria | Sabellaria spinulosa | 0 | 0 | 0 | 0 | 0 | 0 | 327 | 8398 | 0 |
| Sagittoidea | Aphragmophora | Sagittidae | Sagitta | Sagitta elegans | 112 | 511 | 5769 | 10037 | 21403 | 1507 | 1323 | 163 | 0 |
| Sagittoidea | Aphragmophora | Sagittidae | Sagitta | Sagitta setosa | 0 | 0 | 0 | 0 | 0 | 0 | 1007 | 36 | 538 |
| Gastropoda | Cephalaspidea | Scaphandridae | Scaphander | Scaphander lignarius | 106 | 0 | 0 | 0 | 0 | 0 | 0 | 0 | 0 |
| Malacostraca | Mysida | Mysidae | Schistomysis | Schistomysis kerville | 0 | 0 | 0 | 0 | 11844 | 0 | 0 | 0 | 0 |
| Copepoda | Calanoida | Scolecitrichidae | Scolecithricella | Scolecithricella minor | 3579 | 1944 | 0 | 0 | 1526 | 0 | 0 | 0 | 0 |
| Polychaeta | Spionida | Spionidae | Scolelepis | Scolelepis bonnieri | 0 | 19 | 0 | 4701 | 5312 | 11875 | 0 | 60 | 0 |
| Polychaeta | Spionida | Spionidae | Scolelepis | Scolelepis korsuni | 0 | 0 | 0 | 0 | 200 | 521 | 0 | 0 | 0 |
| Actinopterygii | Scombriformes | Scombridae | Scomber | Scomber scombrus | 713625 | 430725 | 2017 | 782499 | 355083 | 121 | 313 | 961 | 2977 |
| Thecostraca | Sessilia | Archaeobalanidae | Semibalanus | Semibalanus balanoi | 0 | 0 | 0 | 63 | 0 | 0 | 0 | 0 | 0 |
| Actinopterygii | Pleuronectiformes | Soleidae | Solea | Solea solea | 0 | 0 | 0 | 0 | 0 | 0 | 0 | 0 | 111 |
| Echinoidea | Spatangoida | Spatangidae | Spatangus | Spatangus purpureus | 1807 | 0 | 0 | 0 | 4180 | 32 | 0 | 0 | 0 |
| Polychaeta | Spionida | Spionidae | Spiophanes | Spiophanes kroyeri | 470 | 0 | 0 | 0 | 124 | 0 | 22 | 0 | 0 |
| Actinopterygii | Clupeiformes | Clupeidae | Sprattus | Sprattus sprattus | 0 | 0 | 0 | 0 | 0 | 0 | 18 | 0 | 1233 |
| Polychaeta | Phyllodocida | Sigalionidae | Sthenelais | Sthenelais limicola | 1207 | 0 | 0 | 0 | 0 | 0 | 0 | 0 | 0 |
| Bivalvia | Galeommatida | Lasaeidae | Tellimya | Tellimya ferruginosa | 0 | 0 | 0 | 0 | 0 | 0 | 0 | 78 | 599 |
| Copepoda | Calanoida | Temoridae | Temora | Temora longicornis | 9968 | 5678 | 20759 | 13702 | 42678 | 148373 | 158679 | 46774 | 170022 |
| Thecostraca | Sessilia | Verrucidae | Verruca | Verruca stroemia | 0 | 0 | 8635 | 1454 | 108 | 0 | 0 | 0 | 39 |

**Table S3: Pearson correlation analysis of the four most abundant copepod taxa. Correlation of relative read abundance (metabarcoding) respectively abundance/$m^3$ (Continuous Plankton Recorder, CPR) and latitude was tested. Significant results are highlighted in bold.**

| **Species** | **Metabarcoding** | **CPR** |
|---|---|---|
| *Oithona similis* (metabarcoding)/ *Oithona spp.* (CPR) | R= -0.15, p= 0.69 | R= 0.08, p=0.17 |
| *Microcalanus pusillus* | R= 0.11, p= 0.78 | / |
| *Calanus finmarchicus* | **R= 0.94, p= <0.01** | **R= 0.2, p= <0.01** |
| *Temora longicornis* | R= -0.62, p= 0.07 | **R= -0.12, p= 0.04** |

**Table S4: Pearson correlation analysis of the four most abundant ray-finned fish taxa. Correlation of relative read abundance (metabarcoding) and latitude was tested. Significant results are highlighted in bold.**

| **Species** | **Metabarcoding** |
|---|---|
| Common mackerel (*Scomber scombrus* ) | R= 0.25, p= 0.51 |
| Common dab (*Limanda limanda* ) | R= -0.5, p= 0.17 |
| Common ling (*Molva molva* ) | **R= 0.77, p= 0.015** |
| Scaldfish (*Arnoglossus laterna* ) | R= -0.53, p= 0.14 |

**Table S5: Pearson correlation analysis of the four most abundant polychaete taxa. Correlation of relative read abundance (metabarcoding) and latitude was tested. Significant results are highlighted in bold.**

| **Species** | **Metabarcoding** |
|---|---|
| *Paramphinome jeffreysii* | **R= 0.76, p= 0.018** |
| *Lagis koreni* | R= -0.089, p= 0.82 |
| *Glycera alba* | R= -0.36, p= 0.34 |
| *Magelona mirabilis* | R= -0.62, p= 0.08 |
